# Bushfire alters the gut microbiome in endangered Kangaroo Island echidnas (*Tachyglossus aculeatus multiaculeatus*)

**DOI:** 10.1101/2023.08.27.554545

**Authors:** Tahlia Perry, Amy Lu, Michael W. McKelvey, Peggy D. Rismiller, Frank Grützner

## Abstract

Kangaroo Island experienced extensive bushfires in December 2019 and January 2020, affecting almost half of the island. This impacted several threatened species including the Kangaroo Island echidnas (*Tachyglossus aculeatus multiaculeatus*). Echidnas can often survive fires, however, changes in soil chemistry and food availability raises questions about the impact on echidna gut health and foraging behaviours. Due to local participation in the citizen science project, EchidnaCSI, echidna scats were collected before and after the fires to directly assess how fire affects their gut microbiomes. Metabarcoding of scat microbiota (n = 13) revealed substantial changes in bacterial composition of echidna faeces post-bushfire when compared to samples collected prior to bushfires. Before the fires, echidna gut microbiomes were more variable and contained mostly soil-associated bacteria, whereas post-fire samples shifted to uniform bacterial communities consisting of lactic acid and gut commensal bacteria. Interestingly, changes were observed in scats collected in both burnt and unburnt regions of the island, suggesting echidnas are foraging between these areas. This is the first study to document changes in gut microbiome of any mammal following bushfires. More work is needed to investigate if the gut bacterial communities continue to change as areas recover from fires.

## 1. Introduction

Fire plays an important role in within the Australian ecosystem, however, extreme weather events such as wildfires are becoming more frequent and intense as climate change extends the number of hot and dry days throughout the year (Bowman et al., 2009; Burrows, 2008; Lindenmayer et al., 2020; Nicholls and Lucas, 2007). Australia experienced its most historically devastating bushfire season from July 2019 to February 2020 (named the Black Summer), which was estimated to burn 97,000 km^2^ of land, affecting 832 vertebrate species (Ward et al., 2020). These fires had a huge effect on plant and animal biodiversity, food availability and habitat (Ward et al., 2020). Kangaroo Island is the third largest island in Australia and from December 2019 to January 2020 almost 50% of the 4,405 km^2^ island was affected by fire (Department for Environment and Water, 2020). This resulted in a call to action to assess the effects of bushfires on priority flora and fauna species (31 plant and 23 animal), including the short-beaked echidna (Department of Agriculture, Water & Environment 2020, ‘Kangaroo Island regional bushfire recovery workshop report’).

Echidnas are within the group of the oldest surviving mammals (monotremes) and well-adapted to fire. Depending on substrate, echidnas can dig underground as the fire front passes. In addition they go into torpor (hibernation-like state) until it is safe to re-emerge (Nowack et al., 2016, McKemey et al., 2019). Echidnas intake a significant amount of soil while foraging and were amongst the first animals observed in the burnt areas once the fires had subsided on Kangaroo Island (P. Rismiller, Pers. Comms.). However, due to the changes in soil chemistry and food availability in the burnt areas, this raised questions about the impact on the gut health and foraging behaviours of the Kangaroo Island echidnas, who were placed on an urgent fire-recovery list for threatened species (Department of Agriculture, Water & Environment 2020, ‘Rapid analysis of impacts of the 2019-20 fires on animal species, and prioritisation of species for management response’).

The gut microbiome plays a vital role in health and wellbeing of animals and an increasing number of studies investigate microbiome changes in wild populations to improve conservation efforts (Bird et al., 2019; Chong et al., 2019; McKenzie et al., 2017). Although there is strong evidence that animals’ gut microbiomes are significantly changed by their environment (especially diet and habitat), research into the effects of bushfires on animal microbiomes is currently limited to a few human oral and gut studies (Perera and Perera, 2018, Gillings et al., 2015). There is, however, a better understanding of how soil microbiomes are affected by fire (Certini et al., 2021; Shen et al., 2016). Microbial communities were initially thought to be resilient to fires, however, they are significantly altered following fire and recovery can take over a decade (Dooley and Treseder, 2012; Pressler et al., 2019). As fires are common in the Australian ecosystem, it will be important to assess how the microbial communities respond regarding native wildlife.

The Kangaroo Island echidna (*Tachyglossus aculeatus multiaculeatus*) is the best studied echidna population in Australia and echidnas can be found anywhere on the island (Rismiller, 1999; Rismiller and Grutzner, 2019; Rismiller and McKelvey, 2003); The Echidna Conservation Science Initiative (EchidnaCSI) is a citizen science project in which researchers collaborate with the community to collect echidna sightings and scat samples (Perry et al., 2022a). Strong participation, in particular on Kangaroo Island, has provided us with samples before and after the 2019/2020 fires to assess the effects of the bushfires on the echidna gut microbiome. Here, we provide the first analysis and comparison of gut microbiomes of echidnas before and after bushfires. Our finding of major microbial changes in scats collected in both burnt and unburnt areas after the fires, raises important questions about the health effects of fire on native Australian species.

## 2. Methods

### 2.1 sample collection and processing

Faecal samples from wild echidnas prior to the 2019/2020 bushfires were collected through a collaborative effort with volunteers as a part of the citizen science project: Echidna Conservation Science Initiative (EchidnaCSI; (Perry et al., 2022a). Two participants had collected echidna scats (n=6) from their properties on Kangaroo Island from October 2017 – October 2019 (Figure 1; Table 1). Faecal samples from wild echidnas after the bushfires were collected by Dr Peggy Rismiller, either found defecated in their habitat or harvested from the lower intestine of a deceased (due to roadkill) echidna (n=7). Roadkill echidnas used in this study were collected by Dr Rismiller under a research permit provided by the Department for Environment and Water, South Australia; Permit number: K16725-32. Two scat samples were collected within the burnt region of Kangaroo Island, while five samples were collected outside the burnt region, on the eastern side of the island (Figure 1; Table 1). After collection, samples were shipped immediately to The University of Adelaide and then stored in a -80 degree freezer.

**Table 1:** Scat samples used in analysis. Fire = whether the scat was collected before or after the 2019 bushfires; fire area = whether the sample was collected in an area affected by fire (Figure 1); sex was known if the scat was taken from a deceased animal and could be correctly identified; echidna breeding season is between June – September.

| Sample ID | Collection Date | Latitude | Longitude | Fire | Fire Area | Material Type | Sex | Season | Breeding Season |
| --- | --- | --- | --- | --- | --- | --- | --- | --- | --- |
| 171002SA1 | 2/10/17 | -35.60 | 137.58 | before | no | scat | unknown | spring | no |
| 171002SA2 | 2/10/17 | -35.60 | 137.58 | before | no | scat | unknown | spring | no |
| 171008SA4 | 8/10/17 | -35.74 | 137.64 | before | no | scat | unknown | spring | no |
| 171205SA1 | 5/12/17 | -35.74 | 137.64 | before | no | scat | unknown | summer | no |
| 180914SA1 | 14/9/18 | -35.74 | 137.64 | before | no | scat | unknown | spring | no |
| 191015SA1 | 15/10/19 | -35.60 | 137.58 | before | no | scat | unknown | spring | no |
| 200125SA1 | 25/1/20 | -35.79 | 137.04 | after | yes | scat | unknown | summer | no |
| 200423SA1 | 23/4/20 | -35.82 | 137.56 | after | no | scat from intestine | male | autumn | no |
| 200508SA1 | 8/5/20 | -35.81 | 137.79 | after | no | scat | unknown | autumn | no |
| 200707SA1 | 7/7/20 | -35.78 | 137.54 | after | no | scat from intestine | male | winter | yes |
| 200707SA3 | 7/7/20 | -35.78 | 137.54 | after | no | scat from intestine | male | winter | yes |
| 200805SA1 | 5/8/20 | -35.77 | 137.49 | after | no | scat from intestine | male | winter | no |
| 200814SA1 | 14/8/20 | -35.99 | 137.04 | after | yes | scat from intestine | male | winter | no |

**Figure 1.**
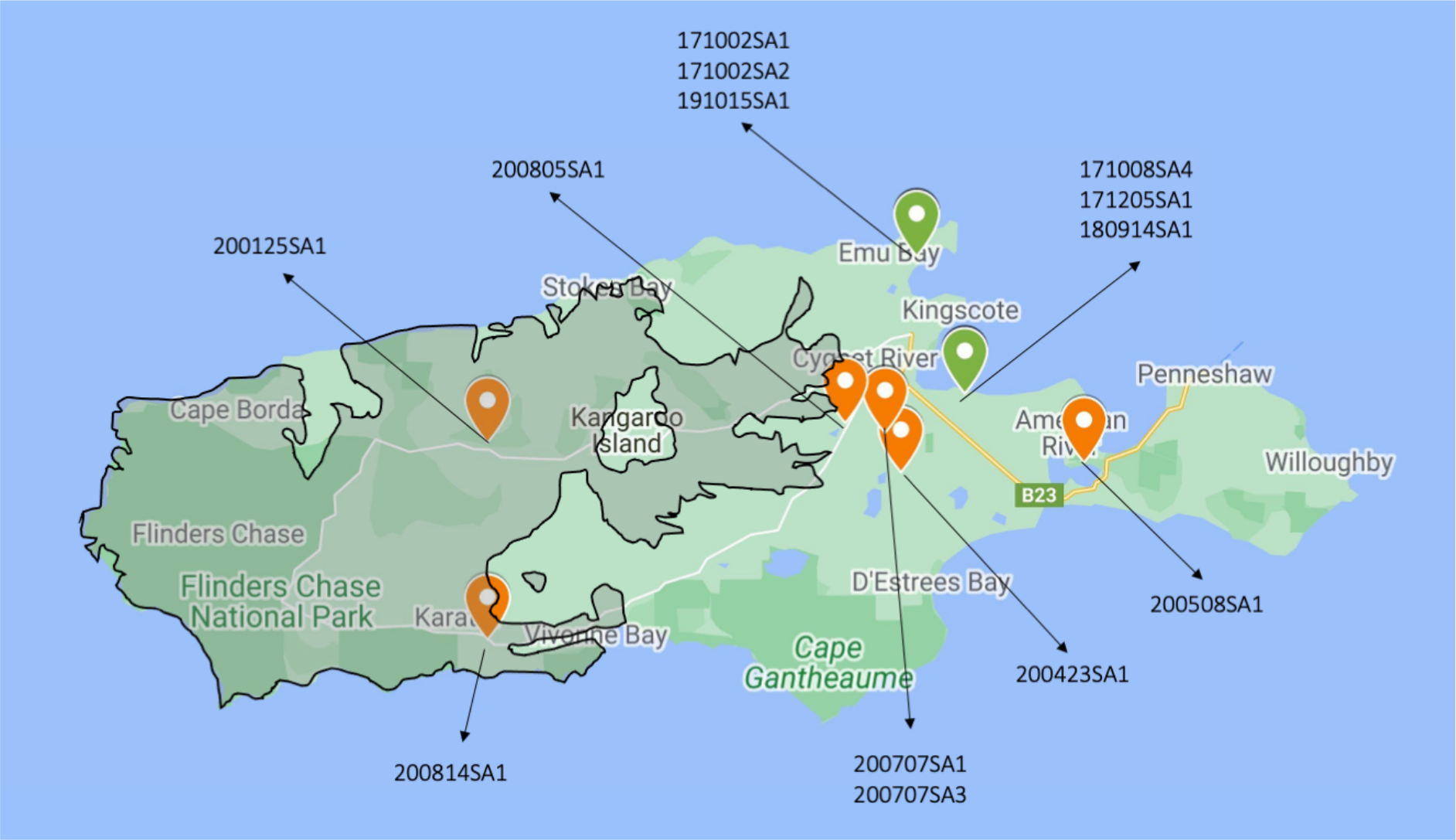
Locations of scat samples collected from Kangaroo Island, South Australia. Samples are labelled with ID number (Table 1); green = samples collected prior to bushfire, orange = samples collected after bushfire. Bushfire impacted regions are shaded grey with black outline.

### 2.1 DNA extraction

Total genomic DNA was extracted from 13 faecal samples using the Qiagen QIAamp Mini Stool Kit (Qiagen, Hilden, Germany) as per the manufacture’s protocol with some modifications. The extractions were performed in a Flow Cabinet Biological Safe Level 2 that was cleaned with 10% bleach (sodium hypochlorite). Approximately a third of the sample was crushed up in liquid nitrogen using a mortar and pestle, prior to adding the sample to InhibitX Buffer. Samples were centrifuged at 20,000 g for 3 min and ∼1 mL eluate transferred to a new 1.5 mL tube. Samples were again centrifuged at 20,000 g for 1 min and ∼700 uL of eluate transferred to a new 1.5 mL tube. Samples were centrifuged one last time at 20,000 g for 1 min and 600 uL added to 25 uL Proteinase K and processed according to the manufacturer’s protocol.

### 2.3 PCR amplification

All samples were PCR amplified and uniquely barcoded, using primers targeting the V4 region of the bacteria 16S ribosomal RNA (rRNA) gene (Caporaso et al., 2011). DNA was amplified with the primer pair 515F (5’-AATGATACGGCGACCACCGAGATCTACACTATGGTAATTGTGTGCCAGCMGCCGCGGTAA-3’) and uniquely barcoded 806R (5’-CAAGCAGAAGACGGCATACGAGATnnnnnnnnnnnnAGTCAGTCAGCCGGACTACHV

GGGTWTCTAAT-3’; Integrated DNA Technnologies). Single reactions of 18.7 μL dH2O, 2.5 μL 10X HiFi buffer, 1 μL 50mM MgSO4, 0.1 μL Platinum Taq DNA Polymerase (ThermoFisher), 0.2 μL 100 mM dNTP, mix, 0.5 μL of 10 μM forward primer, 1 μL of 5 μM reverse primer and 1 μL DNA. DNA was amplified using an initial denaturation at 94 °C for 3 min, followed by 35 cycles of denaturation at 94 °C for 45 sec, annealing at 50 °C for 1 min, elongation at 68 °C for 90 sec, with final adenylation for 10 min at 68 °C, as per accepted Earth Microbiome Protocol (Thompson et al., 2017).

Gel electrophoresis was carried out for each PCR reaction on a 2.5% agarose gel to ensure the samples contained library constructs of desired length (∼390 bp). Each sample was then quantified using Qubit 2.0 Fluorometer and pooled to equimolar concentration. Pooled samples were cleaned following the Agencourt AMPure XP PCR purification protocol (Beckman Coulter), quantified and quality checked by the LabChip® GX Touch™ nucleic acid analyser. A final concentration of 4 nM was run on the Illumina Miseq (v2, 2 x 250bp) at ACRF (Australian Cancer Research Foundation) Cancer Genomics Facility.

### 2.4 Data processing and statistical analyses

DNA sequencing data were processed and analysed using QIIME2 v2022.2 (Bolyen et al., 2019). Demultiplexed paired-end sequence reads were merged, quality filtered and denoised into amplicon sequence variants (ASVs) using the *deblur* plugin (Amir et al., 2017) with a trim length of 250 bp. The feature table was rarefied to a depth of 35,000, using the minimum number of sequences per sample for diversity analysis. Representative sequences were assigned taxonomy using the *feature-classifier* plugin (Naïve Bayesian approach) on the pre-trained SILVA (Quast et al., 2013) 138 V4 region classifier (Bokulich et al., 2018), phylogenetic trees were built using FastTree. Alpha diversity was assessed by diversity metrics, including observed ASVs, Faith’s phylogenetic diversity, Shannon’s entropy and Pileou’s evenness, and statistical significance was assessed using the Kruskal-Wallis tests. Beta diversity was assessed by weighted and unweighted UniFrac metrics and visualized by Principal Coordinates Analysis (PCoA), with statistical significance assessed with Permutational Multivariate Analysis of Variance (PERMANOVA) tests, with 999 permutations.

## 3. Results

### 3.1 Changes in microbial community post-bushfire

DNA sequencing of the 13 scat samples providing an output of 2,373,840 reads, with a mean of 182,603 per sample, resulting in a total of 1,148 Amplicon Sequence Variants (ASVs).

First, we analysed the alpha diversity between samples collected before and after the bushfires, as well as those collected within or outside the burned regions. This revealed that samples collected after the bushfires appeared to have elevated (yet not significant) alpha diversity in comparison to samples collected before the bushfires (p > 0.05; Figure 2). Next, we assessed the microbial compositions of the samples. This revealed a significant change in microbial community for all samples (except one) collected after the bushfires. This was independent of whether they were collected within or outside bushfire affected areas (p = 0.005 unweighted UniFrac; p = 0.011 weighted UniFrac; Figure 3). No effect was observed when comparing seasonal differences or whether the echidna was in breeding season or not (p > 0.05, unweighted and weighted UniFrac).

**Figure 2.**
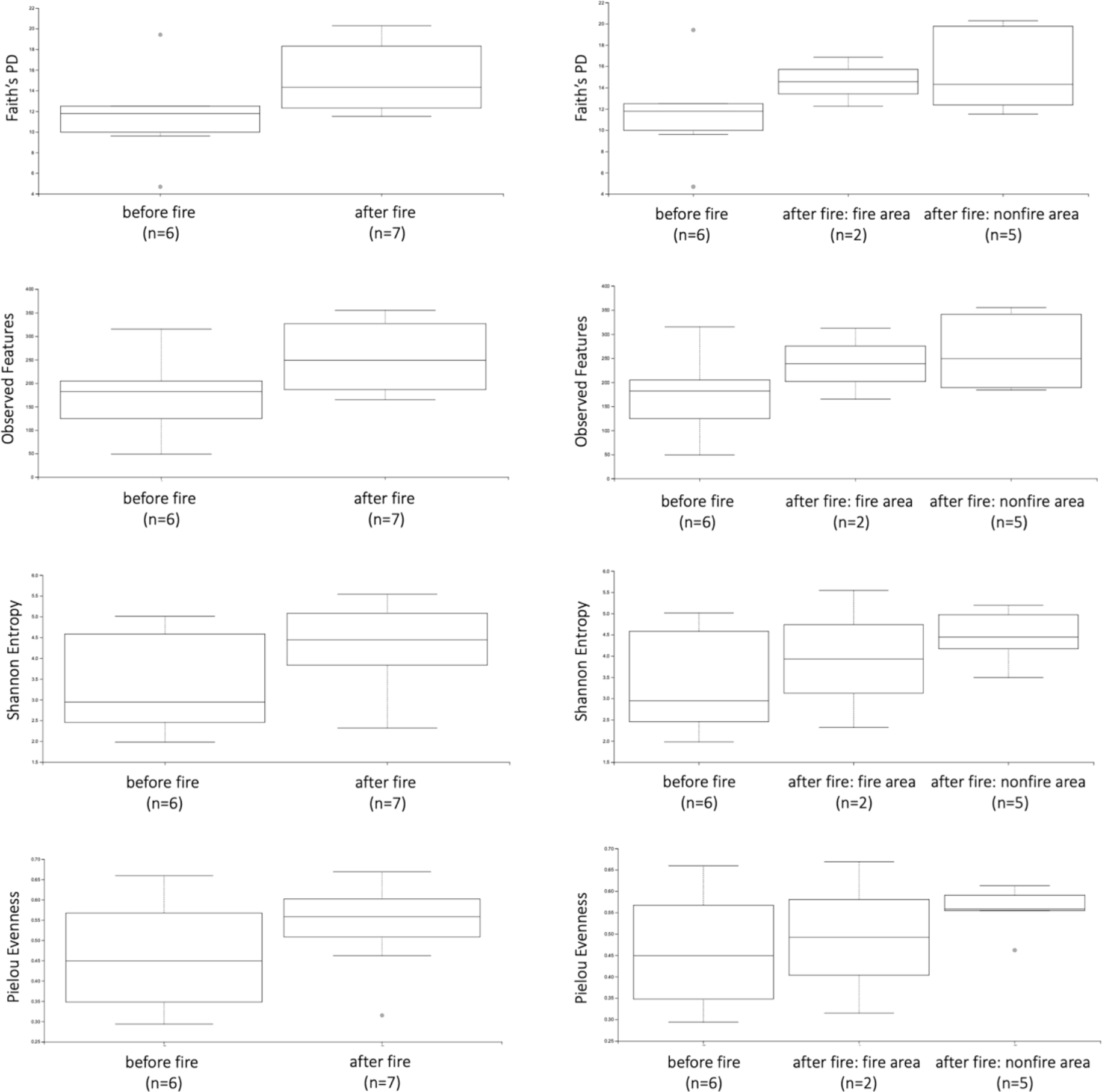
Alpha diversity analyses of scat samples, showing greater microbial diversity in samples post-bushfire. Whisker-box plots depict the following diversity tests: Faith’s phylogenetic diversity (Faith’s PD), Observed features, Shannon’s entropy, and Pielou’s evenness. Left column figures compare if samples were collected the 2019 bushfires to samples collected after the fires, irrespective of region, while the right column figures breaks down the samples collected after the fires either within fire-affected regions (fire area) or outside fire-affected regions (nonfire area; Figure 1). Whisker-box plot horizontal lines indicate median values, upper and lower bounds represent the 25th and 75th percentiles, and top and bottom whiskers indicate maximum and minimum values; outliers are represented with single points. No statistical significance (p < 0.05) was identified between any of the comparisons.

**Figure 3.**
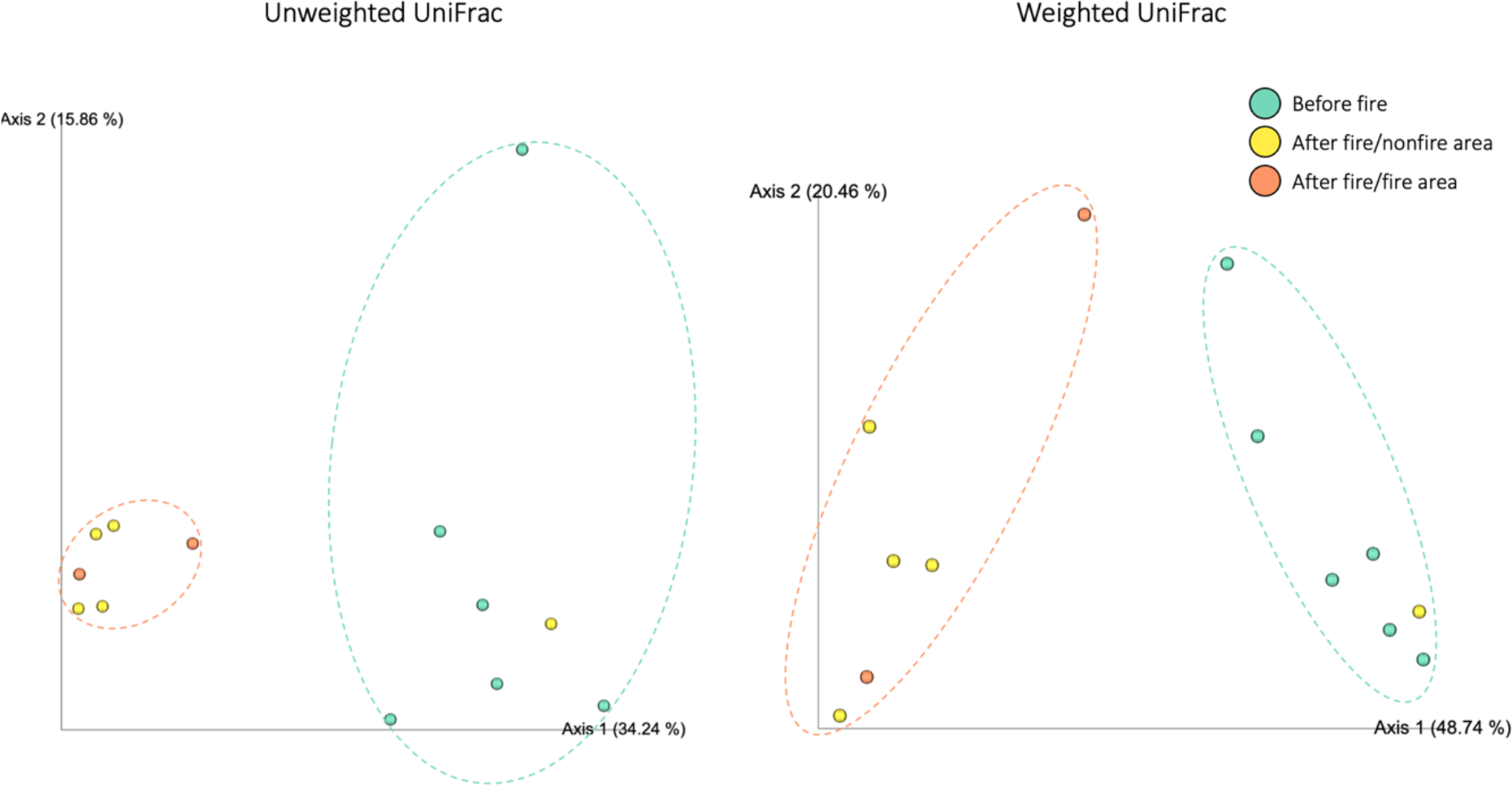
Effect of bushfire on echidna scat microbial composition. Principal Coordinates Analysis plots show unweighted and weighted UniFrac distances for echidna faecal samples. Before fire = sample collected prior to the 2019 bushfires; After fire/nonfire area = after the bushfire but not in a fire-affected region; After fire/fire area = after the bushfire and collected within a fire-affected region. The PCoA plots clearly separate almost all samples collected after the bushfire (dotted orange circle) from samples collected before the bushfire (dotted green circle).

### 3.2 Bacterial taxa changes in samples pre- and post-fire

Visualisation of the bacterial communities within samples, shows that samples collected prior to the bushfires were dominated by bacteria from phyla Proteobacteria, followed by Actinobacteriota and Firmicutes; while samples collected after the bushfires were dominated by Firmicutes, followed by Bacteroidota and Proteobacteria (Figure 4A).

**Figure 4.**
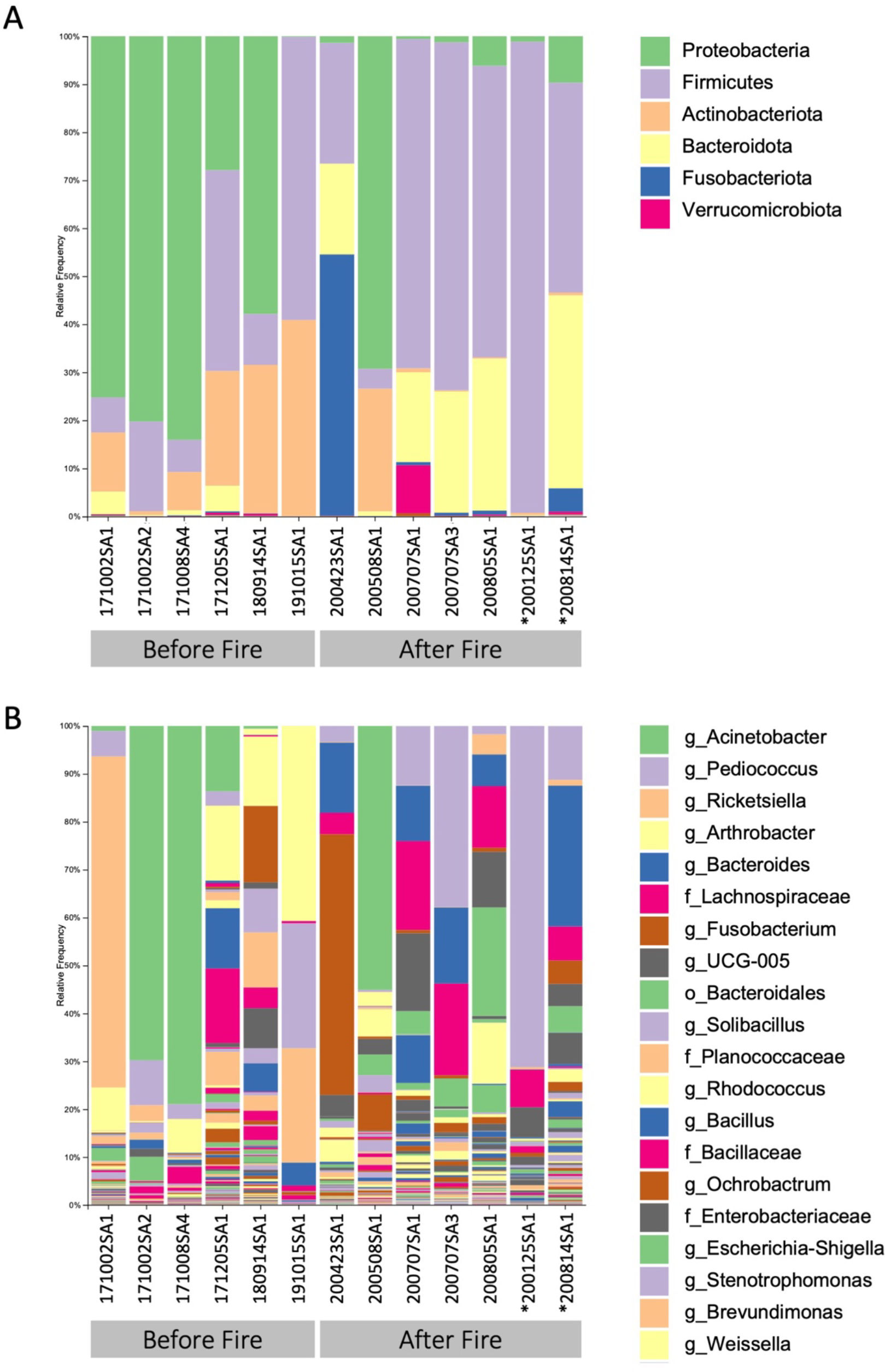
Taxonomy bar plots visualising the effect of bushfire on presence and relative frequency of bacteria in echidna scat samples. A) shows the dominant bacterial phyla present in samples; B) shows top 20 most abundant bacteria genera present in samples. Samples are labelled by their sample ID (Table 1). Before fire = samples collected prior to the 2019 bushfires; After fire = samples collected after the bushfires; * = samples collected in fire affected area; o = order; f = family; g = genus.

At a genus level, there were few genera that were consistently seen in all samples collected prior to the fires (Figure 4B); *Arthrobacter* and *Acinetobacter* were the most common, found in 6/6 and 5/6 samples respectively, with 171002SA2 and 171008SA4 dominated by *Acinetobacter* (70% and 79%, respectively). 171002SA1 was dominated by *Rickettsiella* (69%); 191015SA1 had large proportions of *Arthrobacter* (41%), *Solibacillus* (26%) and unknown genera of *Planococcaceae* (24%); 171205SA1 consisted of relatively even frequency of *Acinetobacter* (14%), *Arthrobacter* (16%), *Bacillus* (16%), and unknown genera of *Bacillaceae* (16%), and was the only sample with a large proportion of *Massilia* (7%). 180914SA1 had the most unique microbial composition with dominating genera consisting of *Rhodococcus* (15%), *Ochrobactrum* (16%), *Stenotrophomonas* (9%), *Brevundimonas* (12%), *Pseudomonas* (4%), *Achromobacter* (8%), and *Patulibacter* (6%).

Samples collected after the bushfires had more similar microbial composition to one another (Figure 4B); The following genera were present in either all or 6/7 samples: *Pediococcus* (0.4– 71%), *Bacteroides* (0.07–29%), *UCG-005* (0.05–16.3%), *Enterobacteriaceae* (0.1–6.5%), *Lactococcus* (0.1–1.5%), *Parabacteroides* (0.6–2.6%), and an unknown genus of *Oscillospiraceae* (0.06–1.6%). There was also a large proportion of an uncharacterised genera of *Lachnospiraceae* (4.5–19%) and several characterised genera belonging to the same family, including *Lachnoclostridium, Roseburia, Tyzzerella*, and *Frisingicoccus. Fusobacterium* was in high proportion (54%) in sample 200423SA1, while in smaller proportions (<5%) across 5 other samples. *Rickettsiella* was seen more consistently in samples collected after bushfires, however, in low frequencies (0.06–4%). 200508SA1 had a similar microbial profile to samples collected before the bushfire; it was mostly dominated by *Acinetobacter* (55%) and the only sample collected after fires to consist of *Arthrobacter* (3%). This sample was collected much further east on the island than the other samples, as seen in Figure 1.

## 4. Discussion

It is well established that the gut microbiome can be affected by environmental changes (Spor et al., 2011). In the context of bushfires, factors impacting on gut microbiota can be related to changes in diet and compositional changes in soil, as well as addition of chemicals used to put out fires. Echidnas are known to survive fires and forage in the affected areas immediately following a fire (McKemey et al., 2019; Nowack et al., 2016). This was confirmed on Kangaroo Island as participants of the EchidnaCSI citizen science project submitted sightings of echidnas immediately following the fires (Figure S1).

Here, we investigated how bushfires affect the gut microbiome of the endangered Kangaroo Island short-beaked echidna. The results show that bushfires significantly influence gut microbiome, where microbial communities in samples collected from echidnas before and after the 2019/2020 bushfires had changed. Interestingly, this was also observed for samples collected outside the burnt areas on Kangaroo Island, which shared the same microbiome signature changes as those collected inside the burnt regions. Echidnas on Kangaroo Island have a home range of up to 88 hectares (Rismiller and Mckelvey, 1994), therefore, it is likely that the scat samples collected outside the burnt regions belonged to echidnas that had foraged within the burnt regions prior to defecating, which is possible as echidnas only defecate once every two days (Snipes et al., 2002). The only post-bushfire sample that had a microbial composition similar to samples collected prior to the bushfires was collected much farther east on the island, where that echidna had most likely not foraged in the burnt areas or was otherwise less exposed to the effects of the fires.

Soil-bacteria were abundant in samples collected before the fires, including *Acinetobacter* (Proteobacteria) and *Arthrobacter* (Actinobacteriota) (Acer et al., 2020; Radkov et al., 2016). This has been documented in echidnas across many different regions in Australia (Perry et al., 2022b). Interestingly the post-fire samples shifted to having more gut commensal (Bacteroidota) and plant-fermenting lactic acid (Firmicutes) bacteria. Soil makes up a significant part of echidna scats, as they ingest soil while foraging. However, intense bushfires will remove a large proportion of topsoil (0-10 cm) as well as change the properties of soil including pH, nutrient content, organic matter content, and the bacterial communities (Ngole-Jeme, 2019; Shen et al., 2016). Echidnas foraging in fire-affected regions are likely not consuming the same bacteria from topsoil as they usually do. Instead, the soil they are ingesting may have a different and more diverse bacterial community, which would explain the greater bacterial diversity seen in scat samples post-fire.

Echidnas are known to eat a variety of insects, worms, beetles and fungi (Feuerherdt et al., 2005; Griffiths, 1968; Rismiller and Grutzner, 2019; Smith et al., 1989; Sprent and Nicol, 2016), with previous microbiome research suggesting that plants form a significant portion of echidna diet (Perry et al., 2022b). Bushfires alter the habitat and food sources available, however, ants and termites are very good at surviving fires (Avitabile et al., 2015; York, 2000). The high abundances of Firmicutes and Bacteroidota in post-fire samples is much more similar to what was observed in the gut microbiomes of eutherian obligate ant and termite eating mammals (Delsuc et al., 2014). Potentially, echidnas foraging in burnt habitat are consuming more ants and termites, and not the varied diet they usually do, which may explain why their microbiomes are more uniform, in terms of the types of bacteria present across samples, in comparison to echidnas feeding prior to the bushfires.

There is surprisingly little research on the effects of fires on gut microbiomes, including a small number on humans (Gillings et al., 2015; Perera and Perera, 2018). As far as we know this is the first study to investigate how bushfires impact gut microbiomes in a non-human mammal. Through EchidnaCSI and local expertise we had the opportunity to investigate this on a small set of samples, which revealed significant changes in echidna microbiomes between samples collected after bushfires on Kangaroo Island compared to those collected prior to the fires. The changes are possibly due to the availability of differing soil composition and food sources. The changes observed in this study encourage more comprehensive future monitoring to assess whether microbial communities continue to change as habitat and food sources recover. Ideally, monitoring will include tracking individuals to determine if echidnas are foraging across burnt and unburnt areas and investigating how quickly the gut bacterial communities change in response to their feeding behaviour and to link this to the overall health of the echidnas. This research highlights that fires can dramatically change gut microbiomes, which raises questions as to the health effects of those changes, not just for echidnas but all wildlife.

## Funding & Acknowledgements

Authors would like to acknowledge the citizen scientists of the EchidnaCSI project, in particular Peter Hastwell and Bill Jenner, who collected echidna scat samples used in this study. This research was funded by the Foundation for National Parks & Wildlife. Tahlia Perry was funded by the Research Training Program. Peggy Rismiller and Michael McKelvey acknowledge the Pelican Lagoon Research Centre for research support and resources. Funding sources had no involvement in study design; in the collection, analysis and interpretation of data; in the writing of the report; or in the decision to submit the article for publication.

## Author Contributions

TP performed sample processing, laboratory work, data analysis, created figures and wrote the manuscript; AL aided in laboratory work, data analysis and figure creation; MWM collected samples and guided the research; PDR collected samples, guided the design of the project, aided in interpretation of results and edited manuscript; FG contributed to design and supervision of the project as well as manuscript preparation and evaluation.

## Data Accessibility

All genetic sequencing data generated for this study are available in the NCBI Sequence Read Archive (SRA) database with BioProject accession number PRJNA952820. All QIIME2 code and outputs can be found at: https://github.com/tahliajperry/2023_SBE_bushfire_microbiome.git

## Ethical Statement

Roadkill echidnas used in this study were collected by Dr Rismiller under a wildlife research permit provided by the Department for Environment and Water, South Australia; Permit number: K16725-32. No ethics approvals are required for using roadkill or faecal material as per the University of Adelaide’s Animal Ethics Committee.

