## Supplementary Figure 1 for "Bushfire alters the gut microbiome in endangered Kangaroo Island echidnas (*Tachyglossus aculeatus multiaculeatus*)"

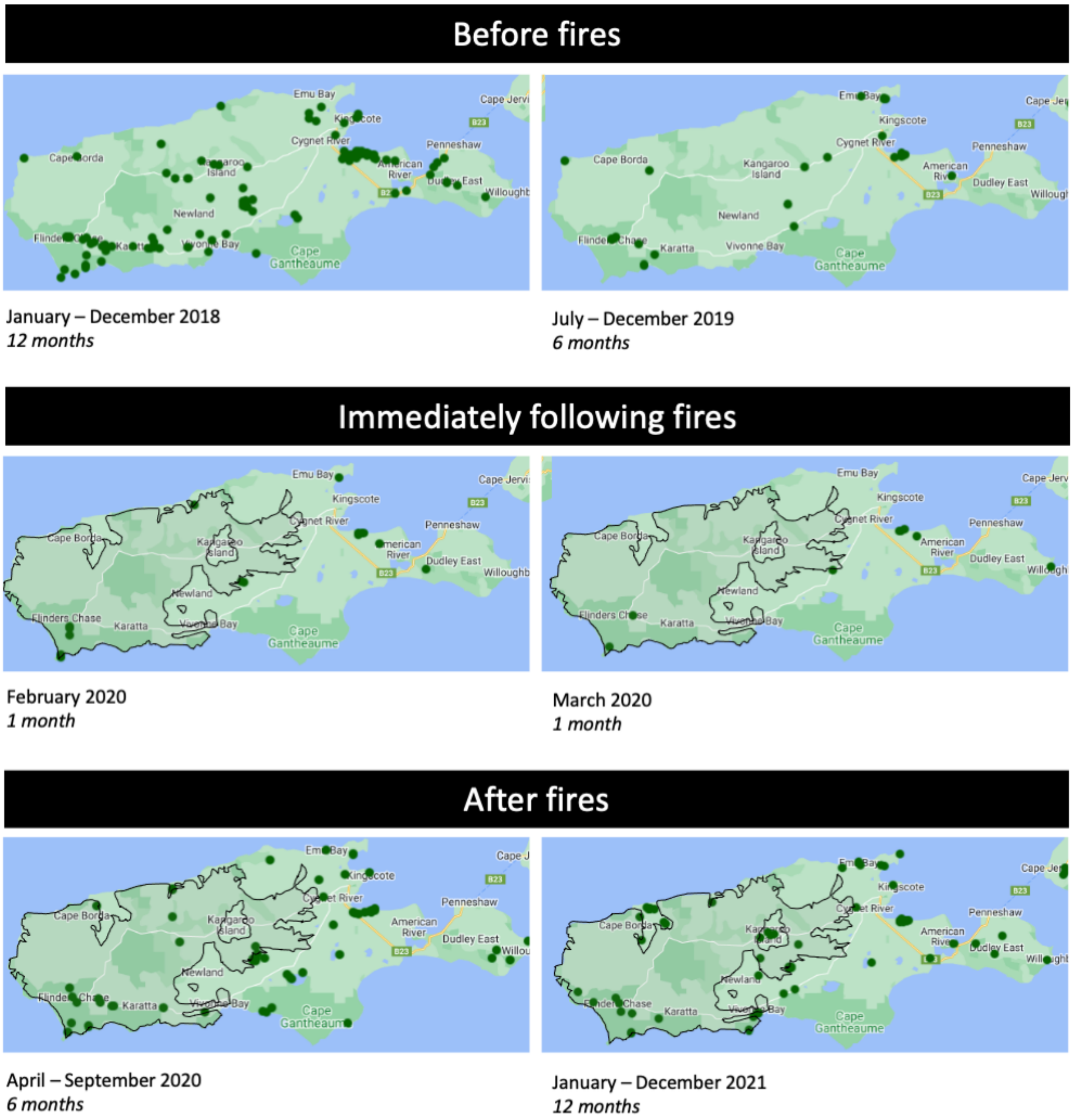


**Figure S1: Sighting data of echidnas across Kangaroo Island (KI).** Bushfire impacted areas of the island are shown with a black outline for fires that occurred in December 2019 and January 2020. Echidnas were seen within the burnt areas of the island immediately following the fires (February and March 2020). As a comparison of the typical number and location of echidnas seen across KI, echidna sightings are also visualised over a 6 and 12 month period before and after the fires. Sighting data was submitted through the national citizen science project, EchidnaCSI. Data is publicly available and was visualised through the Atlas of Living Australia’s Spatial Portal: https//spatial.ala.org.au/layers.
